# Pediatric Severe Sepsis Prediction Using Machine Learning

**DOI:** 10.1101/223289

**Authors:** Thomas Desautels, Jana Hoffman, Christopher Barton, Qingqing Mao, Melissa Jay, Jacob Calvert, Ritankar Das

## Abstract

Early detection of pediatric severe sepsis is necessary in order to administer effective treatment. In this study, we assessed the efficacy of a machine-learning-based prediction algorithm applied to electronic healthcare record (EHR) data for the prediction of severe sepsis onset. The resulting prediction performance was compared with the Pediatric Logistic Organ Dysfunction score (PELOD-2) and pediatric Systemic Inflammatory Response Syndrome score (SIRS) using cross-validation and pairwise *t*-tests. EHR data were collected from a retrospective set of de-identified pediatric inpatient and emergency encounters drawn from the University of California San Francisco (UCSF) Medical Center, with encounter dates between June 2011 and March 2016. Patients (n = 11,127) were 2-17 years of age and 103 [0.93%] were labeled severely septic. In four-fold cross-validation evaluations, the machine learning algorithm achieved an AUROC of 0.912 for discrimination between severely septic and control pediatric patients at onset and AUROC of 0.727 four hours before onset. Under the same measure, the prediction algorithm also significantly outperformed PELOD-2 (*p* < 0.05) and SIRS (*p* < 0.05) in the prediction of severe sepsis four hours before onset. This machine learning algorithm has the potential to deliver high-performance severe sepsis detection and prediction for pediatric inpatients.

## Introduction

Sepsis is a high-impact condition that affects both adults and children. In 2001, the total burden of sepsis-spectrum syndromes in the United States was estimated at $16.7 billion and 215,000 deaths annually.^1^ In 2007, the mean, per-hospitalization cost of severe sepsis was estimated to be $47,126,^2^ and a recent study assessed that sepsis is responsible for as many as 5.3 million deaths per year globally.^3^ Pediatric sepsis in particular causes over 6,500 deaths annually in the United States, with an estimated $4.8 billion burden of care, at approximately $64,280 per hospitalization.^4^ Moreover, survivors can suffer both short-term^5^ and long-lasting impacts.^6^

Relative to that of adult sepsis, the literature of pediatric sepsis is less developed. This includes consensus definitions of pediatric sepsis,^7 8 9^ which may not match clinicians’ diagnoses in practice.^10^ The current pediatric consensus definitions for sepsis^6^ are closest to the older, three-stage adult sepsis definitions,^11^ rather than the more recent, two-stage “Sepsis-3” definitions.^12^

As is true for adult sepsis,^13 14^ many studies show that early and aggressive treatment of pediatric sepsis with antibiotics and fluids correlates with better outcomes.^15 16 17 18 19^ While there is some evidence that fluid bolus treatment is detrimental in pediatric patients,^20^ it is nevertheless true that determining the most effective treatment depends on early detection and diagnosis. Traditionally, the call for early detection has been answered by severity scoring systems, which are lacking in specificity for pediatric sepsis. For example, while the Systemic Inflammatory Response Syndrome (SIRS) criteria have been adapted for pediatric patients,^7^ they are intended to assess infection and other inflammatory response in general. In some cases, scoring systems for nonspecific pediatric disease severity or mortality are applied to the task of recognizing pediatric sepsis, such as the Pediatric Logistic Organ Dysfunction score (PELOD-2).^21^ In the absence of specialized pediatric sepsis scores, some hospitals have implemented home-grown computerized sepsis prediction systems, which may benefit from site specificity.^22^

Computerized prediction systems offer a compelling alternative to manual application of scoring systems. Such systems access electronic health record (EHR) data to inform recommendations to clinicians. These systems have the potential to catch septic patients who might otherwise be missed, typically while in the emergency department, and could provide early warning of sepsis for patients in intensive care. Studies in adults show that this latter setting of hospital-acquired sepsis among inpatients is both distinct and substantially more deadly.^23 24 25^ Tools such as the Modified Early Warning Score (MEWS)^26^ or other sets of rules can be used to rank adult patients by some measure of their risk of developing sepsis.^27 28 29^ The scoring systems on which these approaches are based, however, tend to be created and evaluated on large cohorts (e.g., qSOFA^30^) without regard to special, site- or population-specific conditions, which might render them less effective than site-specific prediction tools.

Machine-learning (ML)-based approaches can be easily customized using site- or population-specific data, ultimately resulting in improved performance relative to generic scoring systems^31 32 33^ and to non-customized applications of the same ML-based system. ^34 35 36^ While ML-based systems have been applied to prediction of sepsis in neonatal patients, conditioned on the availability of real-time waveform data^37 38^ or extensive sets of laboratory and historical data,^39^ they have not previously been applied to EHR-based prediction for the pediatric inpatient population. If successful, such predictors could provide easily-accessible, site-customized early sepsis warning for pediatric patients. In the experiments discussed below, our objective was to create and demonstrate such a customized, high-performance ML-based prediction tool for pediatric severe sepsis.

## Methods

### Data Set

In these experiments, we use de-identified chart data from pediatric (ages 2 to 17 years) inpatient and emergency encounters at the University of California San Francisco (UCSF) Medical Center, from June 2011 to March 2016, inclusive.^40^ This age range corresponds to the older three pediatric subpopulations as described by Goldstein.^7^ The original UCSF data collection did not impact patient safety and all data were deidentified in accordance with the Health Insurance Portability and Accountability Act (HIPAA) Privacy Rule prior to commencement of this study. Hence, this study constitutes non-human subjects research which does not require Institutional Review Board approval. The data set includes 11,621 encounters. After exclusion, described below, the final data set includes 11,127 encounters, of which 103 are labeled as severely septic. The inclusion flowchart and demographic characteristics of the data set are presented in Figure 1 and Table 1.

**Figure 1:**
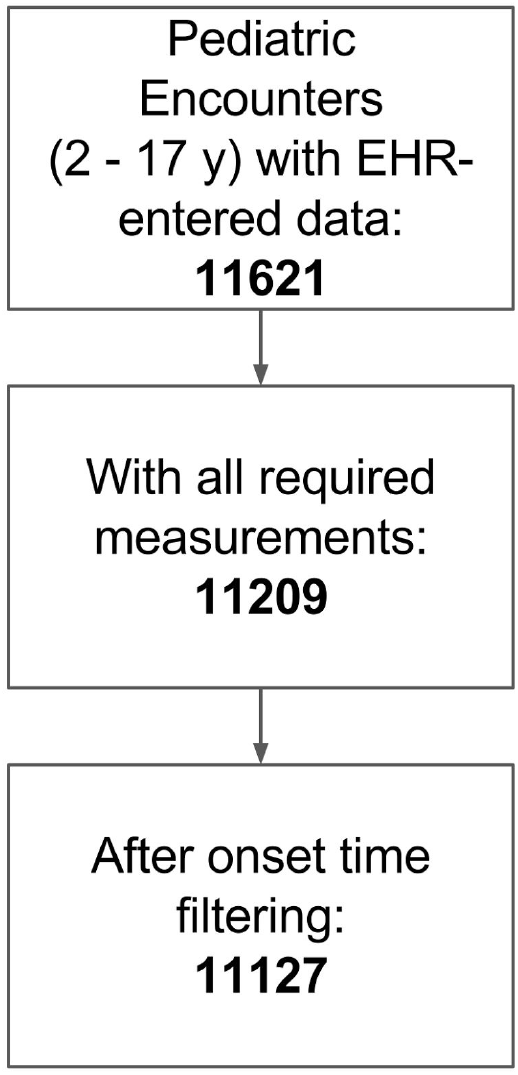
Inclusion flowchart. The final data set used in training and testing constitutes 11,127 examples, of which 103 (0.93%) are labeled as severely septic.

**Table 1:** Demographic information of pediatric inpatients at UCSF from June 2011 to March 2016, inclusive.

| Characteristic |  | Overall |  | Severe Sepsis |  |
| --- | --- | --- | --- | --- | --- |
|  |  | Count | Percent | Count | Percent |
| Gender | Female | 5,494 | (49.38%) | 46 | (44.66%) |
|  | Male | 5,633 | (50.62%) | 57 | (55.34%) |
| Age<br>Overall:<br>Median 10,<br>IQR (5-14)<br>Severe sepsis:<br>Median 11,<br>IQR (4-14) | 2-5 | 3,101 | (27.87%) | 34 | (33.01%) |
|  | 6-12 | 4,081 | (36.68%) | 26 | (25.24%) |
|  | 13-17 | 3,945 | (35.45%) | 43 | (41.75%) |
| Length of Stay<br>(days)<br>Overall:<br>Median 3, IQR<br>(1-6)<br>Severe sepsis:<br>Median 21,<br>IQR (10-46) | 0-2 | 4,919 | (44.21%) | 1 | (0.97%) |
|  | 3-5 | 3,344 | (30.05%) | 7 | (6.80%) |
|  | 6-8 | 1,189 | (10.69%) | 11 | (10.68%) |
|  | 9-11 | 536 | (4.82%) | 17 | (16.50%) |
|  | 12+ | 1,073 | (9.64%) | 66 | (64.08%) |
|  | Unknown | 66 | (0.59%) | 1 | (0.97%) |
| In-Hospital<br>Death | Yes | 67 | (0.60%) | 16 | (15.53%) |
|  | No | 11,060 | (99.40%) | 87 | (84.47%) |

### Data Processing and Screening

The UCSF EHR data were organized into a SQL database and custom queries were used to extract the vital sign, lab report, and other data used in our experiments. These patient records were loaded by Dascena’s proprietary software to prepare examples for training and prediction.

Encounters were removed if the recorded patient age was less than two or more than eighteen years; this range corresponds to non-infant age brackets in the consensus sepsis definitions.^7^ In addition, encounters were removed if they were missing any of the required measurements (patient age, diastolic and systolic blood pressures, heart rate, temperature, respiration rate, and peripheral oxygen saturation) to be used in training and prediction; while supplemental measurements (Glasgow Coma Score, white blood cell count, and platelet count) were passed to the training and testing routines, their presence was not required. The inclusion diagram is presented in Figure 1. From 11,621 encounters with appropriate ages, 11,209 remained after checking for the required measurements. As a final step, encounters with severe sepsis onset too early in the stay (< 6 hours after the start of the patient record) were removed, such that the set of encounters used at varying pre-onset offsets (see Experimental Procedures) was constant. Encounters with any other error in onset time determination were also removed. After removing encounters for these onset times, a total of 11,127 encounters remained for training and testing. Of these examples, 103 (0.93%) were labeled severely septic.

For encounters meeting the inclusion criteria, observations of vital signs were binned by the hour and simple carry-forward imputation was used when no observation was available for a given hour. Based on these observation time series, we constructed a variety of derived features (e.g., approximate Mean Arterial Pressure constructed as a linear combination of systolic and diastolic blood pressures) and calculated the sepsis gold standard (see Supplemental Tables 1 & 2).

### Gold Standard

The gold standard follows the pediatric severe sepsis definition of Goldstein et al., 2005, wherein severe sepsis requires:

- SIRS score of ≥ 2, where at least one of temperature or white blood cell count is abnormal;
- suspicion of infection, operationalized here as the presence of an ICD-9 code for septicemia, sepsis, severe sepsis, or septic shock, which might be attached at any time during the encounter (given that the patient meets the SIRS criteria above, this is “sepsis” under the Goldstein criteria); and
- Organ dysfunction

Under the Goldstein criteria, septic shock is further defined by when the above conditions are met and there is *cardiovascular* organ dysfunction. Pediatric SIRS criteria, the gold standard, and the organ dysfunction criteria (part of the gold standard) are presented in Supplemental Tables 1, 2, and 3, respectively.

The gold standard was implemented by electronic chart abstraction, combining data entered into the EHR throughout the encounter with ICD-9 codes. All of these criteria follow Goldstein et al., 2005,^7^ with modifications necessary for application to the UCSF pediatric data set. These necessary modifications include those allowing for binary white blood cell count (normal vs. abnormal), lack of radiological information, and lack of patient history. Without information on physician intention and examination observations, fluid administration in the presence of low blood pressure was assumed to be an attempt to resuscitate, and a section of the Goldstein cardiovascular dysfunction definitions was removed (see Supplemental Tables for details of implemented criteria).

Our gold standard is particularly concerned with the timing of severe sepsis onset. Timing information comes from the satisfaction of the SIRS criteria and the organ dysfunction criteria. When the patient first meets the SIRS criteria is the time of sepsis onset (if a sepsis-related ICD-9 code, which does not carry timing information, is present). When a patient who, at some point in their stay, meets the SIRS criteria and has a sepsis-related ICD-9 code (is “septic” under our gold standard), this patient is marked as having severe sepsis with an onset time defined by the first time the organ dysfunction criterion is met. Note that this means that our retrospective definition may determine that a patient is “severely septic” before they meet the surveillance criteria for being “septic.” This feature was deemed necessary both to reflect the clinical process and to avoid difficulties surrounding noisy satisfaction of the thresholds.

### Modeling

All learning conducted in this work was done using boosted ensembles of decision trees.^41 42^ Ensemble classifiers combine the output from many “weak” learners, each of which would be insufficient to solve the desired learning problem on its own, creating a strong learner. These decision trees each carry out a sequence of multiple binary classifications, where each is of the form of a measurement value compared against a threshold. The sequence of comparisons and the required thresholds for each are created during the training process. The appropriate set of branching checks is performed for each tree within the final classifier, traveling along the tree structure until a leaf node (and corresponding risk score) are reached. The risk scores from the individual trees are then aggregated to assign an overall risk score.

Our classifiers were trained on a set of features that included patient age, diastolic and systolic blood pressures, heart rate, temperature, respiration rate, and peripheral oxygen saturation (SpO_2_). As noted above, encounters had to have all of these measurements at some point during their stay to qualify for inclusion in the analyses. Additionally, the values of Glasgow Coma Score, white blood cell count, and platelet count were used if available. The final feature vectors were organized, along with their gold standard labels, into arrays to be passed to the training and prediction routines.

### Experimental Procedures

We compared the performance of the algorithmic sepsis predictor with that of the concurrent, running values of the PELOD-2 and pediatric SIRS scores. These experiments used all patients of at least 2 and less than 18 years of age in the data set, treated as one aggregate population. This population was split into four approximately equal-sized “folds” (sets) for four-fold cross-validation (CV).^42^ The CV procedure allows the estimation of generalization performance and its variability, as well as comparison of this performance with the PELOD-2 and SIRS scores, calculated hourly. Due to the original data set’s encoding of laboratory values as only normal/abnormal, the affected subscores of PELOD-2 were approximated with 1 point for abnormal and 0 points for normal. We computed a variety of metrics on the performance of the resulting classifiers (and the PELOD-2 and SIRS scores) on the test folds. We determined statistical significance using one-tailed paired *t*-tests, where each pair constituted the same metric of two different classification methods, measured on the same test fold. The p-value threshold for significance was fixed at 0.05 for all comparisons. We repeated these experiments for pre-onset offsets of 0, 1, 2, 3, and 4 hours, examining our system’s ability to learn pre-onset patterns in septic patients.

### Data Availability

Any inquiries regarding the dataset used in this study can be addressed to the corresponding author.

## Results

We evaluated predictive performance of the ML-based predictor by training and testing at hourly intervals from sepsis onset and through four hours before onset. Figures 2 and 3 show the performance of the algorithm at two such times in terms of ROC curves, which show the tradeoff between sensitivity (the number of severe sepsis examples detected over all severe sepsis examples) and specificity (the number of false alarms given, over all examples that are not severely septic). In comparing two ROC curves, a superior classifier is one that has either better sensitivity for a fixed specificity (is higher on the plot) or better specificity for a fixed sensitivity (is farther to the left). In the onset time discrimination plot, the ML-based predictor’s ROC curve is nearly strictly dominant over the PELOD-2 and SIRS curves, and has a larger area under it (i.e., larger AUROC). Figure 4 shows how cross-validation-fold-averaged AUROC varies as a function of prediction horizon in hours for each prediction system. These comparisons are statistically significant (*p* < 0.05, one-tailed pairwise *t*-test) for 1 and 4 hours pre-onset (PELOD-2) and 0, 1, and 4 hours pre-onset (SIRS). Table 2 presents a set of detailed performance metrics for the algorithm, SIRS, and PELOD-2. Apart from AUROC, these performance metrics are a function of a chosen operating point (i.e. a point on the ROC curve).

**Figure 2:**
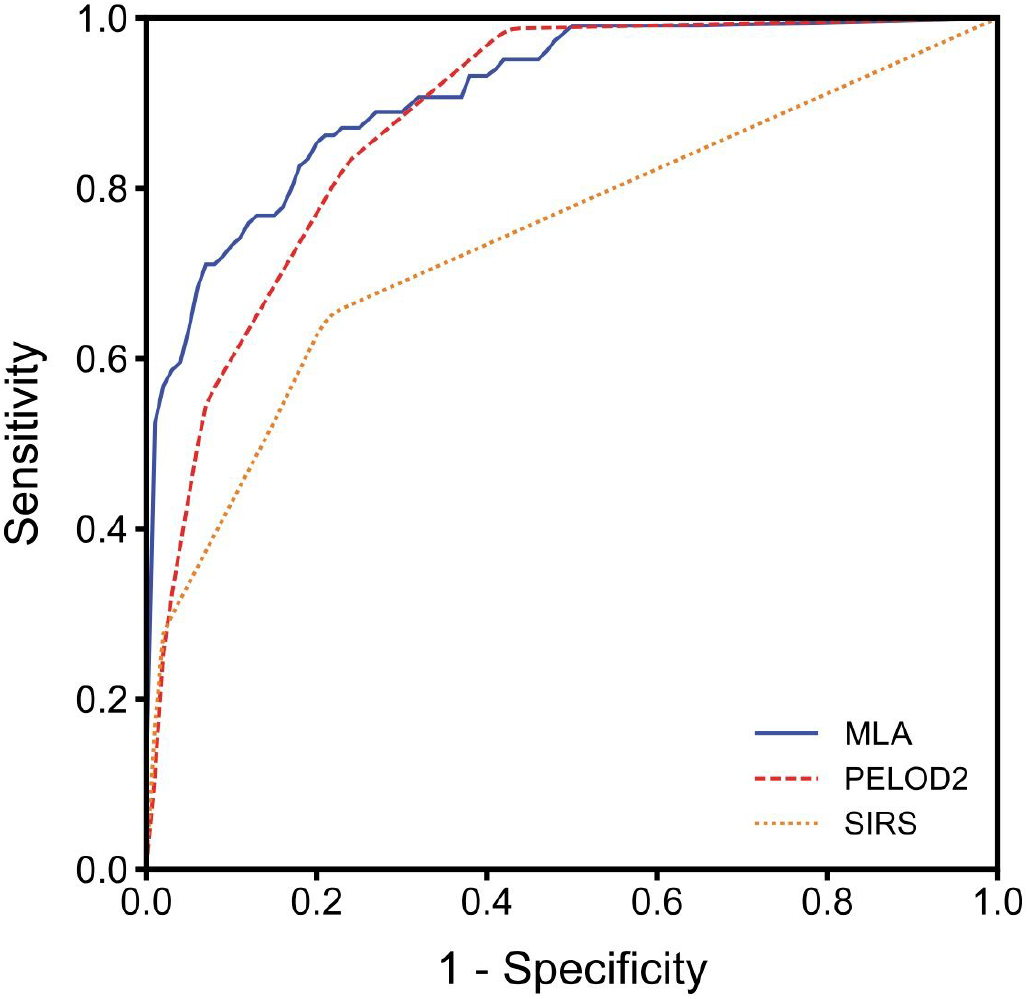
ROC curves (averaged across the four test folds) for the machine learning algorithm (MLA), PELOD-2, and SIRS at time of onset.

**Figure 3:**
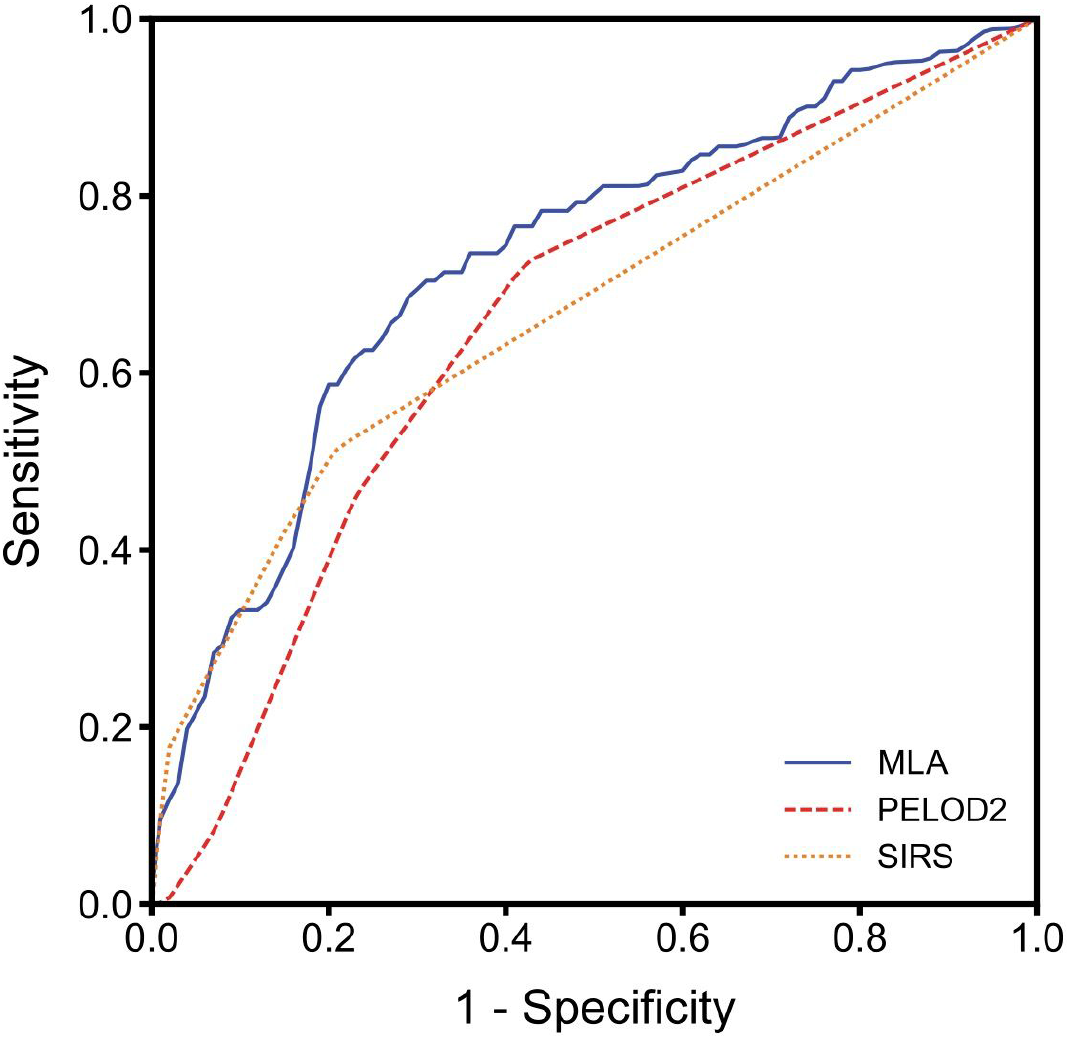
ROC curves (averaged across the four test folds) for the machine learning algorithm (MLA), PELOD-2, and SIRS at 4 hours pre-onset.

**Figure 4:**
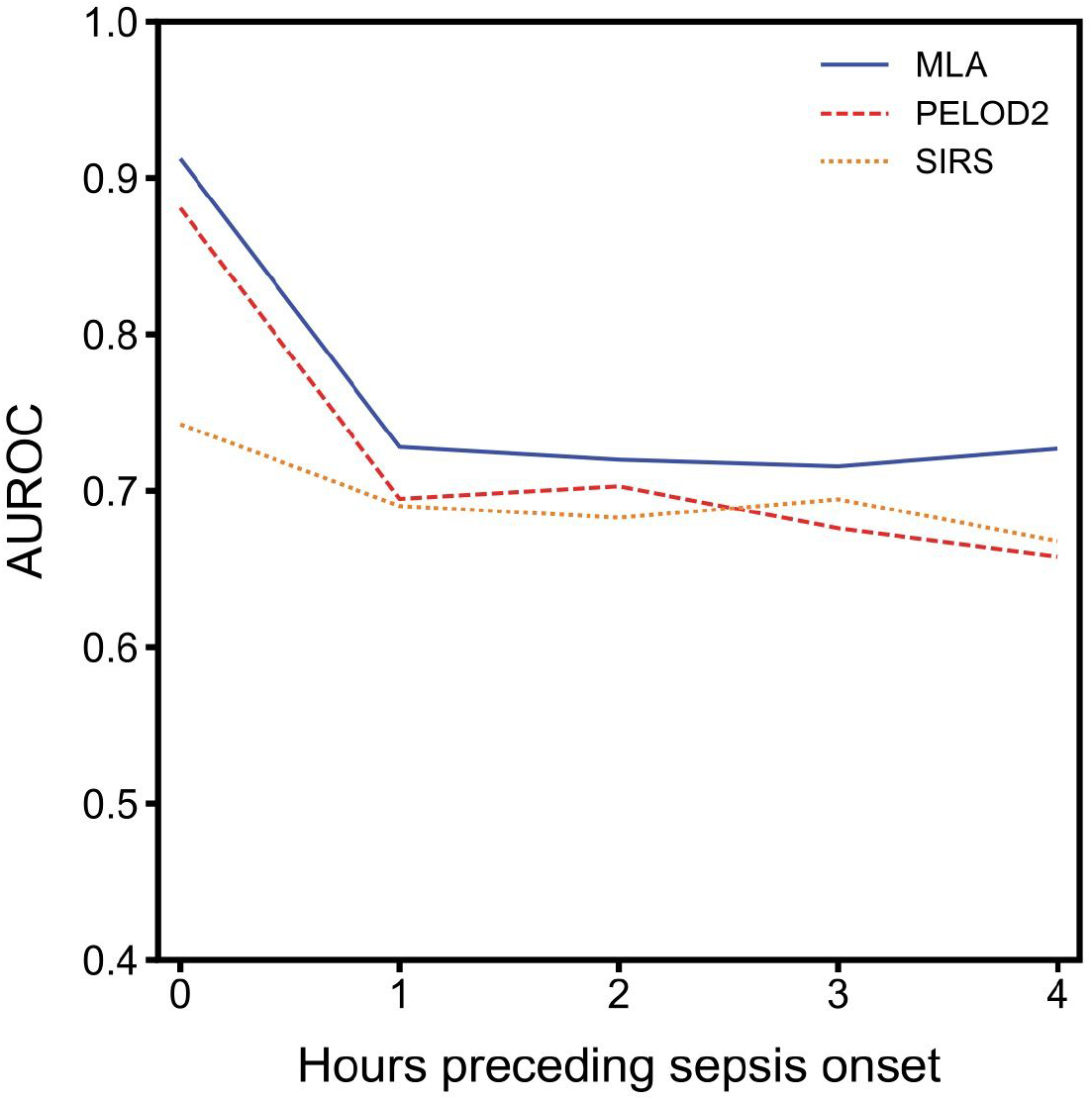
Average AUROC over a prediction horizon. These AUROC differences are statistically significant for the machine learning algorithm (MLA) versus PELOD-2 at 1 and 4 hours pre-onset (p < 0.05) and versus SIRS at 0, 1, and 4 hours pre-onset (p < 0.05). Non-significant comparisons against PELOD-2 have pvalues of 0.0596, 0.2819, and 0.2270 (0-, 1-, and 3-hour). Nonsignificant comparisons against SIRS have p-values of 0.0652 and 0.3335 (2- and 3-hour).

**Table 2:**
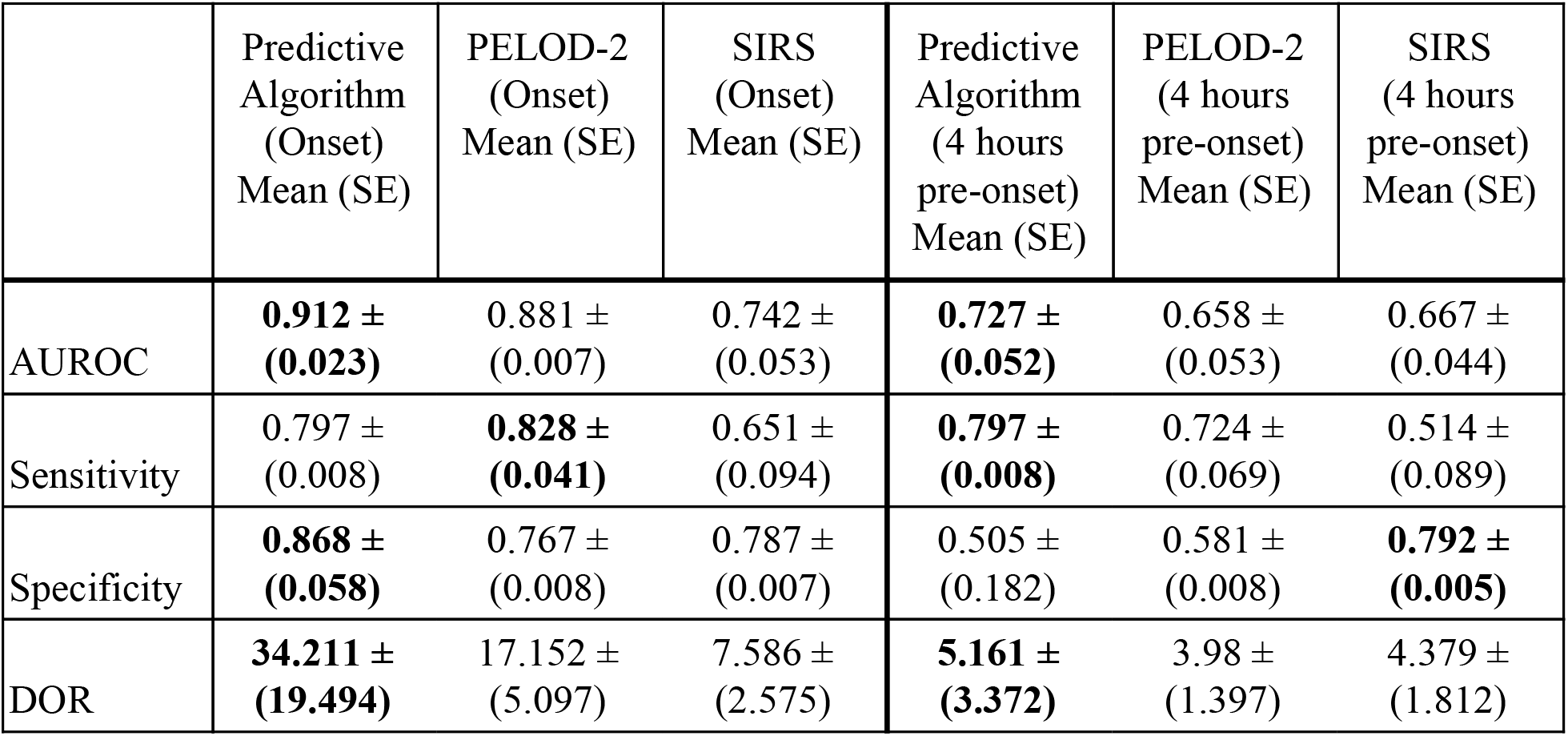
Performance metrics for the machine learning algorithm and pediatric scoring systems. For each metric and each time (onset or 4-hours pre-onset), the best result is bolded. This procedure chose an operating point from the ROC curve where the sensitivity was the largest possible value ≤ 0.80; the selected PELOD-2 and SIRS sensitivity values for 4-hour pre-onset prediction were considerably below this value, allowing them to obtain favorable tradeoffs in some of the other metrics. SE is the standard error and DOR is the diagnostic odds ratio.

|  | Predictive<br>Algorithm<br>(Onset)<br>Mean (SE) | PELOD-2<br>(Onset)<br>Mean (SE) | SIRS<br>(Onset)<br>Mean (SE) | Predictive<br>Algorithm<br>(4 hours<br>pre-onset)<br>Mean (SE) | PELOD-2<br>(4 hours<br>pre-onset)<br>Mean (SE) | SIRS<br>(4 hours<br>pre-onset)<br>Mean (SE) |
| --- | --- | --- | --- | --- | --- | --- |
| AUROC | **0.912** ±<br>(0.023) | 0.881 ±<br>(0.007) | 0.742 ±<br>(0.053) | **0.727** ±<br>(0.052) | 0.658 ±<br>(0.053) | 0.667 ±<br>(0.044) |
| Sensitivity | 0.797 ±<br>(0.008) | **0.828** ±<br>(0.041) | 0.651 ±<br>(0.094) | **0.797** ±<br>(0.008) | 0.724 ±<br>(0.069) | 0.514 ±<br>(0.089) |
| Specificity | **0.868** ±<br>(0.058) | 0.767 ±<br>(0.008) | 0.787 ±<br>(0.007) | 0.505 ±<br>(0.182) | 0.581 ±<br>(0.008) | **0.792** ±<br>(0.005) |
| DOR | **34.211** ±<br>(19.494) | 17.152 ±<br>(5.097) | 7.586 ±<br>(2.575) | **5.161** ±<br>(3.372) | 3.98 ±<br>(1.397) | 4.379 ±<br>(1.812) |

## Discussion

These experiments demonstrate that the ML-based sepsis prediction system can predict severe sepsis onset with AUROC performance superior to that of existing pediatric organ dysfunction and inflammatory response scoring systems (Table 2, Figure 2, Figure 3 & Figure 4). These comparisons were statistically significant versus PELOD-2 (organ dysfunction) at 1 and 4 hours pre-onset and versus SIRS at 0, 1, and 4 hours pre-onset. This superiority is also visible in other metrics, particularly at onset time.

The ML-based system can be a competitive and useful means of assessing pediatric patients’ likelihood of developing severe sepsis. This clinical problem represents a significant opportunity for clinical decision support, as it is critical to both provide monitoring for this particularly vulnerable population and avoid excessive numbers of alarms. Our machine learning algorithm outperforms the PELOD-2 and pediatric SIRS scoring systems, indicating that it has the potential to deliver these essential improvements. While neither PELOD-2 or SIRS are primarily intended for sepsis prediction, they provide a practical baseline for this retrospective study. These can be computed in the absence of biomarkers, nursing reports, or access to other authors’ machine learning-derived models. In this context, these results are particularly compelling in light of the granularity of our continuous, chart-abstracted nature of data set and gold standard, which allow us to determine a sepsis onset time within an inpatient encounter.

The gold standard is a possible limitation in the present analysis. First, chart review would provide a superior gold standard, but it is not applicable at scale, requiring the present use of our surveillance-type gold standard. Second, by limiting “suspicion of infection” to those who have ICD-9 codes for sepsis-spectrum syndromes, we prevent the gold standard from positively labeling encounters not acknowledged as being on this spectrum; this could mean that the gold standard is under-reporting sepsis prevalence, were the definition used in this work to be applied prospectively. Further, Weiss et al.^10^ compared the clinical diagnoses of severe sepsis by attending physicians with the result of the application of the Goldstein consensus definitions and found that the agreement between the two was only moderate (Cohen’s *κ*, 0.57 ± 0.02, mean ± SE). On a more technical note, the current analysis uses only ICD-9 codes and does not use ICD-10 codes, while the study period includes the roll-out of the ICD-10-CM coding system and the mandatory compliance date of October 1, 2015.^43^ However, ICD-9 codes for sepsis appear with a similar frequency both before and after the roll-out date. Finally, while the gold standard gives a particular onset time, it is difficult to assess how this relates to severe sepsis onset as would be observed by an attending clinician.

The characteristics of the UCSF pediatric inpatient population may limit generalizability; these data are from a tertiary care center with a particularly heavy representation of organ transplant patients. This population also has a low prevalence of hospital-acquired severe sepsis (< 1%), limiting the power of the statistical analyses.

For future, more complex work in pediatric sepsis prediction via machine learning, an important requirement is obtaining other and larger data sets. Data sets with higher sepsis prevalence, whether due to the population served or some other factor, might be particularly helpful in this regard. Likely candidates for obtaining such data sets are other tertiary care facilities. It should be noted that, while pediatric sepsis is generally rare, even secondary-care contexts could benefit from being able to identify the few cases that do appear, particularly if such capabilities could be integrated into a larger data collection and prediction ecosystem, as they can with this predictive algorithm.

In summary, the ML-based sepsis prediction system examined in these experiments outperforms traditional, tabular scoring systems and demonstrates superior performance in predicting pediatric severe sepsis onset. The improved ROC performance offers clinicians and hospitals a variety of useful operating points to suit their sepsis alerting needs. The ROC performance also offers the promise of using the actual numerical score produced by this algorithm for severe sepsis risk stratification. Using these tools, clinicians will be better able to allocate finite clinical resources, catch patients before their condition deteriorates, and avoid adverse outcomes. This work also represents a novel application of machine learning techniques for sepsis detection in the pediatric population. We hope that this contribution encourages others to tackle this intriguing and challenging problem.

## Acknowledgements

The authors gratefully acknowledge Dr. Julie Fitzgerald for her insightful suggestions on the manuscript, and Anna Lynn-Palevsky and Emily Huynh for suggestions and help with editing.

## Additional Information

### Author Contributions

All authors listed on this manuscript were involved in the conception and design of this study, with TD and QM responsible in particular for the development of the machine learning algorithm. All authors had the opportunity to draft and revise this manuscript, and all have approved it in this final form. All authors agree to be accountable for the integrity and accuracy of the contents of this paper.

## List of Abbreviations

AUROC: area under the receiver operating characteristic curve
CV: cross-validation
DOR: diagnostic odds ratio
EHR: electronic health record
ICD-9: international classification of diseases, 9th revision
IQR: interquartile range
ML: machine learning
MLA: machine learning algorithm
PELOD: Pediatric Logistic Organ Dysfunction score
ROC: receiver operating characteristic
SE: standard error
SIRS: Systemic Inflammatory Response Syndrome
UCSF: University of California San Francisco

